## Supplemental for "Structure of the IL-27 quaternary receptor signaling complex"

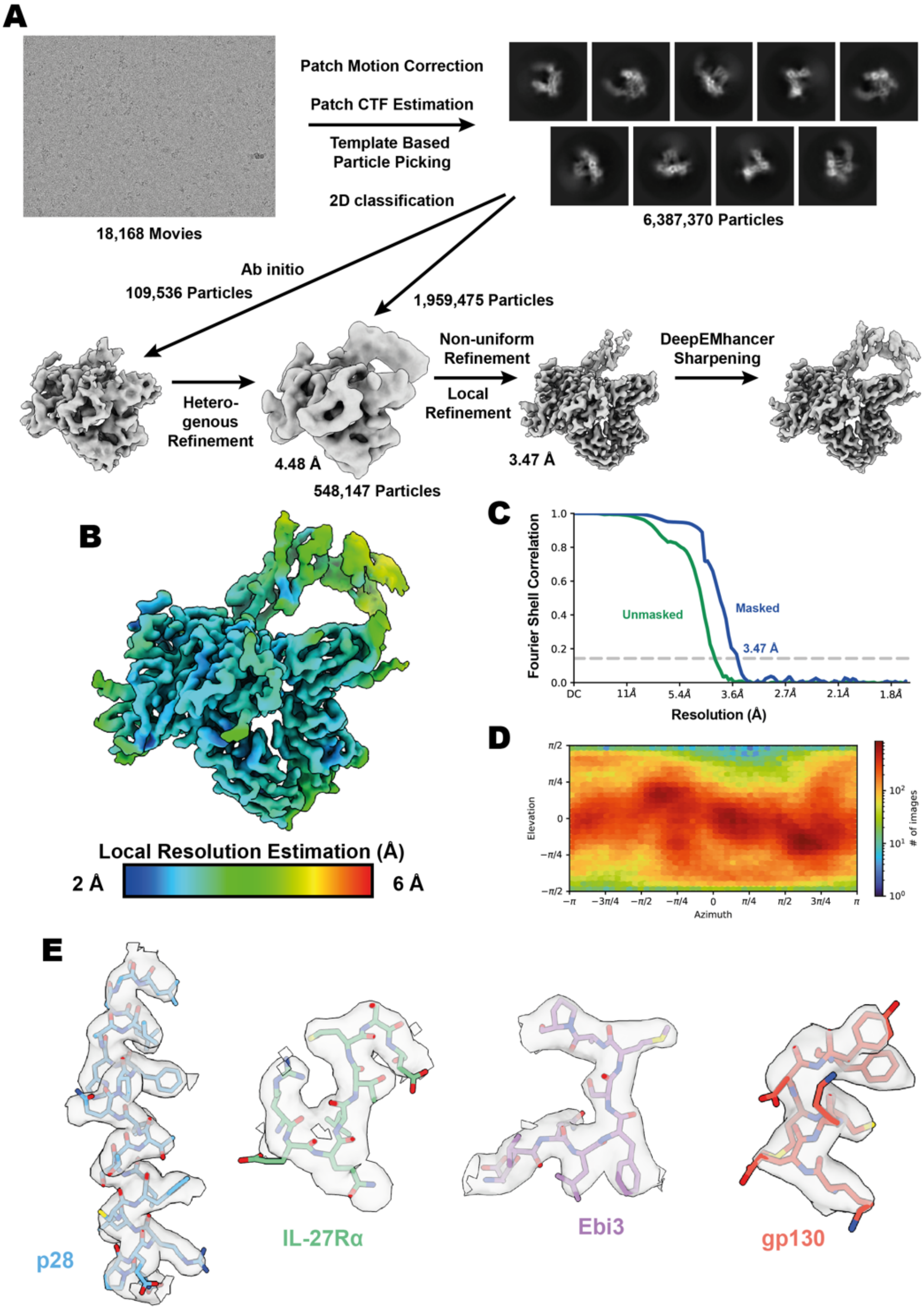

**Supplemental Figure 1. IL-27 quaternary complex cryoEM data processing.** (**A**) Workflow for cryoEM data processing. Representative micrograph, reference free 2D averages, and cryoEM maps at the various stages of processing. Depicted maps were z-flipped prior to model building. (**B**) Local resolution estimation of the finalised cryoEM map of IL-27 quaternary complex (Punjani et al., 2017). (**C**) FSC curve of the IL-27 quaternary complex reconstruction using gold-standard refinement calculated from unmasked and masked half maps. (**D**) Orientational distribution of the IL-27 quaternary complex reconstruction. (**E**) Representative cryoEM density regions of p28, IL-27Rα, Ebi3, and gp130.

**Supplementary Table 1. CryoEM data collection, refinement, and validation statistics.**

| | IL-27 Complex (PDB/7U7N 7LQ6/EMD-26382) |
|---|---|
| **Data collection and processing** | |
| Magnification | 105,000 |
| Voltage (keV) | 300 |
| Electron exposure ($e^-/Å^2$) | 60 |
| Defocus range (μm) | -0.8 to -2.0 |
| Pixel size (Å) | 0.839 |
| Symmetry imposed | C1 |
| Initial particle images | 6,387,370 |
| Final particle images | 548,147 |
| Map resolution FSC threshold (Å) | 0.143 |
| Map resolution (Å) | 3.47 |
| **Refinement** | |
| Initial model used (PDB) | AlphaFold |
| Model resolution FSC threshold (Å) | 0.143 |
| Model resolution (Å) | 1.9 |
| Map sharpening *B*-factor ($Å^2$) | 189.6 |
| Model Composition | |
| Non-hydrogen atoms | 7,284 |
| Protein residues | 884 |
| Ligands | 16 |
| *B*-factors ($Å^2$) | |
| Protein | 102.97 |
| Ligand | 114.62 |
| R.m.s. deviations | |
| Bond lengths (Å) | 0.003 |
| Bond angles (°) | 0.610 |
| Validation | |
| MolProbity score | 1.60 |
| Clashscore | 8.81 |
| EMringer score | 2.33 |
| Rotamer outliers (%) | 0.89 |
| Ramachandran plot | |
| Favoured (%) | 97.37 |
| Allowed (%) | 2.63 |
| Outliers (%) | 0.00 |
